# Investigation of astrocytes’ morphological changes in response to laser-induced shockwave

**DOI:** 10.1101/2023.11.29.569124

**Authors:** Pegah Pouladian, Janelle Ho, Nicolas Perez, Nicole M. Wakida, Veronica Gomez-Godinez, Daryl Preece

## Abstract

Traumatic Brain Injury (TBI) arises from an external force affecting the brain, leading to a range of outcomes from mild to severe. Despite continuous scientific advancements, it continues to pose a persistent threat and remains a significant cause of physical impairment and mortality.

Various models, including blast-induced TBI (bTBI), have been proposed to simulate TBI. Laser-induced shockwaves (LIS) us emerging as an effective method. LIS generates shockwaves via pulsed laser-induced plasma formation, offering a controlled means to study TBI at the cellular level. Astrocytes, pivotal in maintaining brain function post-injury, undergo dynamic morphological changes, contributing to the understanding of injury responses and neurodegenerative diseases.

This study introduces a system combining Laser-Induced Shockwaves (LIS) and Quantitative Phase Microscopy (QPM) to quantify morphological changes in astrocytes during and after LIS exposure. QPM, a label-free method, facilitates 3D imaging and captures real-time cellular dynamics. The integration of LIS and QPM enables the assessment of astrocyte responses to shear stress caused by LIS, revealing immediate and sustained morphological transformations.

Analysis post-LIS exposure indicates significant alterations in circularity, volume, surface area, and other features. Statistical tests affirm of observed trends, providing insights into astrocyte responses to mechanical forces. The findings contribute to understanding how mechanical stimuli impact astrocyte morphology, holding promise for targeted therapeutic strategies in traumatic brain injuries and related neurological disorders. The integrated LIS and QPM approach serves as a powerful tool for 3D imaging and quantitative measurement of astrocyte morphological changes, offering deeper insights into cellular dynamics and potential therapeutic interventions.

## Introduction

Traumatic Brain Injury (TBI) stands as a critical public health concern, with a notable surge in TBI-related incidents reported by the Centers for Disease Control and Prevention from 2006 to 2014 [1]. This injury occurs when an external force disrupts the normal function of the brain, leading to a spectrum of outcomes, from mild to severe, with consequences ranging from full neurological recovery to mortality. Despite advancements, TBI remains a leading cause of physical impairment and death, particularly among the youth.

Various models have been proposed to simulate TBI. These models can be categorized as acceleration models of TBI, compression models of TBI, repetitive models of mild TBI, and blast models of TBI. There has been an increase in blast-induced TBI (bTBI) caused by improvised explosive devices impacting the brain directly. Different methods of blast models of TBIs have been proposed [2]. One method in particular, laser-induced shockwaves (LIS), has been shown to be an effective way to simulate shockwaves in vivo [3] and in vitro [4]. Laser-Induced Shockwave (LIS) is a phenomenon that occurs when a fluid is irradiated by pulsed laser light, leading to plasma formation and the creation of a cavitation bubble. This bubble’s expansion causes a shockwave, which can be used to damage cells by mechanical force.

When an injury occurs to the brain, neuronal cells in the brain interact with each other to maintain the brain’s normal function. Astrocytes, the most numerous cells in the CNS, play a crucial role in maintaining equilibrium between ions, maintaining homeostasis of water and blood flow, recycling neurotransmitters, and supplying the nutrition that cells need to remain healthy [5, 6]. They are imperative to the brain’s normal functioning after CNS injury, regulating leukocyte infiltration, repairing the blood–brain barrier (BBB), protecting neurons, and restricting nerve fiber growth [7]. Astrocytes possess complex morphologies in terms of size, shape, and processes. The base cell structure has a star-like shape, with the soma being in the center and the processes protruding in the outward direction. These processes also display smaller branching processes, which also vary in shape and size [8]. The majority of individual astrocyte processes branch out and do not overlap between neighbors, forming the gliapil, or region where thousands of synapses can be embedded. The thin processes approaching these synapses are believed to be important sites for neuron-astrocyte interactions due to their close proximity [9].

The field of astrocyte morphology is experiencing rapid growth, reflecting a heightened interest and increased research efforts aimed at comprehending the structural characteristics of these cells. The dynamic nature of astrocyte processes in vivo, marked by boundary shifts, adds a layer of complexity to their study. However, analyzing the arrangement of astrocytes poses a challenge due to the asynchronous addition of these cells from embryonic to adult stages and the inherently three-dimensional nature of the system. Potential mechanisms contributing to the formation of astrocyte domains include homotypic repulsion, resulting in limited overlap, and an alternative process involving initial overlap followed by competitive ramification until exclusive territories are established [10]. In vivo, astrocytes within their designated territories maintain a consistent volume fraction and an increase in segment density during the early postnatal weeks. This indicates an amplification in astrocyte branching and a growth in the number of smaller yet thinner astrocyte processes throughout this developmental period [9]. Changes in astrocyte morphology have been directly observed in specific brain regions during different neurodegenerative diseases, such as Parkinson’s disease (PD), Huntington’s disease (HD), and amyotrophic lateral sclerosis [11].

The injury and disease response process of astrocytes in the CNS is referred to as astrogliosis. Depending on the severity of the insult and the proximity of the astrocyte to the injury, the degree of cellular hypertrophy is heterogeneous and highly variable among reactive astrocytes. The different forms of morphological heterogeneity of astroglia are also directly correlated with the degree of hypertrophy and the degree of interaction and interdigitation of cell processes [12].

In this study, we introduce a system that enables quantitative phase imaging of astrocyte cells before, during, and after LIS exposure to stimulate TBI at a cellular level. We capture 3D images of the LIS injury and quantitatively measure the changes in the cell membrane and internal cell structure. Quantitative Phase Microscopy (QPM) is an emerging label-free method used to study transparent cells and tissues without photo-bleaching effects often encountered with conventional fluorescence imaging. This modality uses interferometry and precisely quantifies the optical path length caused by the sample, enabling the ability to image transparent features in cells, measure the movements of their organelles, and quantify the cellular dynamics including membrane motility [13–15]. This system allows for the application of different degrees of shockwave injury while monitoring changes in bulk membranes, cell shape, and real-time damage and recovery of intracellular damage. By integration of LIS and QPM, the system is capable of 3D imaging of LIS injury and quantitatively measures membrane changes, as well as internal cell structure in astrocyte cells in response to shear stress caused by LIS. The examination of how astrocytes react to laser-induced shockwaves (LIS) reveals immediate and sustained morphological transformations, providing insights into the dynamic adaptations of these cells. Following LIS, astrocytes swiftly shift to a more circular shape, accompanied by alterations in volume, surface characteristics, and height features. Subsequent analysis conducted 2 hours post-LIS indicates enduring changes, suggesting continuous cellular adjustments and regulatory processes. Statistical analysis underscores significant alterations in circularity, volume, surface area, and other features, highlighting the dynamic responses to mechanical disturbance. Mann-Kendall Tau-b test results affirm the robustness of the observed trends. The dynamics of surface area imply a cellular response geared towards enhancing chemical interactions, while an increased projected area and area-to-volume ratio indicate a flattening response to LIS. The findings from this research illuminate significant morphological changes in astrocytes in response to LIS, contributing to the growing body of knowledge on how mechanical stimuli impact astrocyte morphology [16, 17]. These insights hold promise for further investigations into the underlying mechanisms of astrocyte responses to mechanical forces and the development of targeted therapeutic strategies in the context of traumatic brain injuries and related neurological disorders. The integrated LIS and QPM approach offers a powerful tool for 3D imaging and quantitative measurement of astrocyte morphological changes, providing deeper insights into cellular dynamics and potential therapeutic interventions [16, 17].

## Materials and methods

### Cell preparation

An established astrocyte type-I (Ast-1) line (clone CRL-2541) was received directly from ATCC. Ast-1 cells were cultured in advanced DMEM media supplemented with 2% FBS, 1% glutamax, and 0.2% gentamicin/amphotericin B. Ast-1 cells were plated onto glass-bottom 35mm imaging dishes. Alternatively, imaging dishes were coated by incubating them for 1 hour in a 1:60 dilution of matrigel:DMEM to promote cell adhesion.

### Laser induced shockwave setup

As depicted in Figure 1, for Laser-Induced Shockwaves (LIS) setup, a Q-switched diode-pumped solid-state (DPSS) laser *(Flare NX, Coherent)*. This laser emits pulses with a duration of 1.5 ± 0.2 nanoseconds at a wavelength of 1030 nanometers and can operate at frequencies up to 2000 Hz. To generate laser pulses, the laser is connected to a function generator GFG-8015G (*GW Instek*), producing a 5-volt square wave to trigger the laser pulse generation.

**Fig 1.**
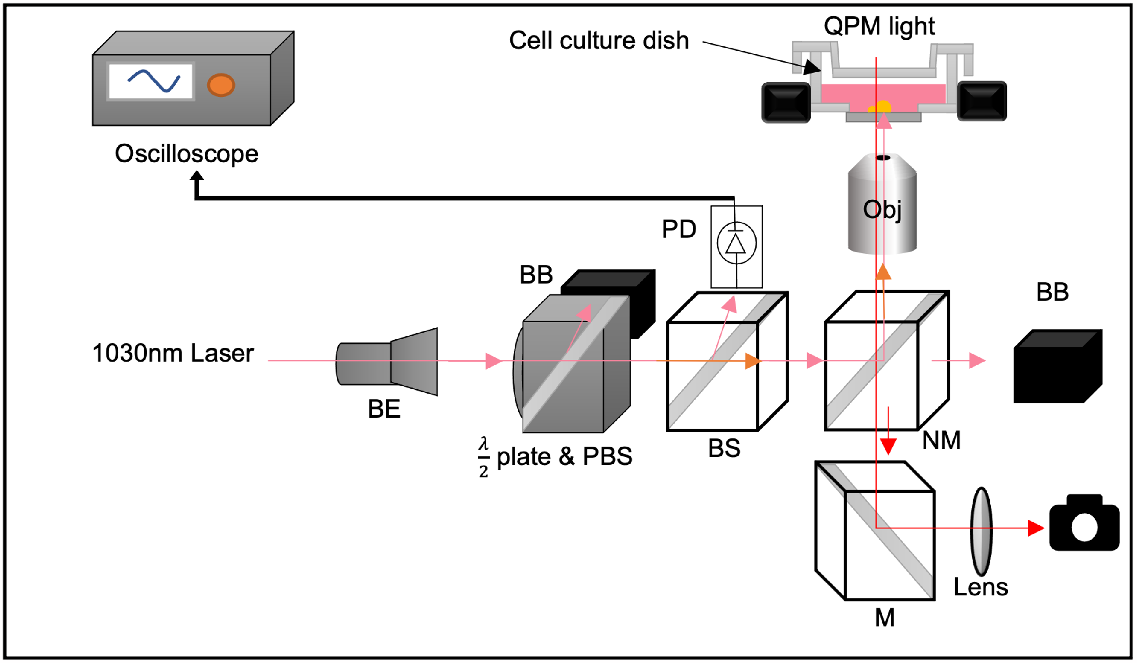
Laser-induced shockwave setup.

After passing through a 3x beam expander, the laser power is controlled by a half-wave plate plorizer and a polarized beam splitter. The remaining portion of the beam is reflected by a near-infrared (NIR) mirror and focused onto the cell culture medium from underneath using an oil immersion objective lens *(Plan-Apochomat 40x/1.3, Zeiss)*.

The laser power is measured at the back aperture of the objective lens with a thermal sensor S425C (*Thorlabs*), connected to the Optical Power and Energy Meter Console PM400 (*Thorlabs*).

## 1 Cell segmentation and feature extraction

The field of cell biology has evolved significantly since the establishment of cell theory in the 19th century, which recognized the cell as the fundamental unit of life [18]. Light microscopy, in particular, has played a pivotal role in advancing cell biology [19], building upon the pioneering work of Antoni van Leeuwenhoek in the 1670s. Presently, cell biologists have access to a diverse array of advanced microscopic imaging techniques that allow for the visualization of cellular phenomena, surpassing the classical diffraction limit of light. These techniques generate extensive and complex datasets that have outgrown manual management, processing, and analysis. Consequently, the application of computational techniques for these tasks has become indispensable for the advancement of cell biology [20–24].

One of the primary challenges in cell studies is image segmentation, a critical step that involves the precise delineation of cell boundaries. Accurate cell delineation is pivotal for subsequent analyses, regardless of whether they focus on nanoscale details or encompass broader millimeter-scale perspectives [25].

Cell segmentation refers to the process of partitioning an image or series of images into distinct regions that correspond to individual cells. The main objective of cell segmentation is to detect and delineate each cell present in an image for further analysis. This step is crucial in image-based research in the field of biology and biomedicine. It is widely used in various biological and biomedical studies, including cell tracking, cell enumeration, investigation of cell behavior, and other forms of cellular analysis. To perform cell segmentation, it is essential to create an accurate mask that outlines the cell boundaries. Two critical factors are taken into account when developing a cell segmentation method: the ability to identify the borders of each cell correctly and the ability to differentiate between adjacent cells. Various methods have been tested and evaluated, and in the following sections, we will discuss some of these approaches in detail.

### 1.1 Approaches for cell segmentation procedure

Various plugins, such as MorphoLibJ, Stardist, and local thresholding followed by Classic Watershed, were tested for different fields of view (FOV) containing astrocyte cells. The study aimed to evaluate their capabilities in identifying astrocyte cells and accurately masking their borders. These image-processing techniques were combined with the ‘regionprops’ function in MATLAB to facilitate the isolation of individual cells for further analysis. Each tested approach was assessed based on its ability to differentiate neighboring astrocyte cells and its effectiveness in generating precise cell masks, particularly with respect to cellular processes.

#### 1.1.1 Watershed

The watershed algorithm is a popular method for segmenting images in computer vision and image processing. This algorithm works particularly well when identifying objects or regions in an image that have clear boundaries. The algorithm mimics a flooding process, and it interprets pixel intensities as topographical elevations. This idea was first introduced by S. Beucher and C. Lantuéjoul [26]. The process starts by placing markers in the image at the highest and lowest intensity points, and these points guide the flooding process, assigning regions with common markers the same label. The output is a segmented image with clear boundaries shown by watershed lines. The watershed algorithm is used in many areas, including medical imaging for cell segmentation and tumor detection, as well as in geological and satellite image analysis.

##### Program Overview

Fiji/ImageJ has a watershed tool that can be accessed in the binary section. The tool works by treating the input image as a topographic surface with water sources placed at regional minima using the conventional watershed model. As the algorithm floods the relief, it creates dams where the water sources converge. However, this program has limited options for adjustments.

#### 1.1.2 MorphoLibJ (Distance Transform Watershed 3D)

##### Program Overview

Distance Transform Watershed 3D is a segmentation tool that is available for free and open-source. It uses a combination of distance transform and watershed segmentation concepts to perform the segmentation. The distance transform feature computes the distance of each voxel to the nearest background voxel. The watershed algorithm utilizes these distance values as the water levels flood the image from local minima until the watershed lines are formed. Users have the option to customize the distance map and the watershed settings, including dynamic and connectivity settings.

#### 1.1.3 Stardist

##### Program Overview

Stardist is a deep learning-based image segmentation tool that can be easily installed as a plugin for Fiji/Image J. It employs a fully convolutional neural network to detect and delineate objects using a unique “star-complex” shape model. This shape model uses a star-convex polygon concept which enhances segmentation accuracy, thanks to the presence of multiple “arms” in the segmentation shape. Additionally, the model utilizes a probability map to further refine the segmentation, estimating the likelihood of each pixel belonging to the object. Finally, a binary mask is generated that identifies the object pixels, and post-processing parameters such as non-maximum suppression (NMS) settings can be adjusted by users, including probability/score and overlap threshold.

Figure 2 presents a sample field of view (FOV) and the outcomes yielded by various methodologies. Despite meticulous parameter tuning to optimize accuracy, none of these techniques exhibited promising results. In Figure 2(c), the classic watershed plugin result is depicted, wherein the algorithm fails to effectively distinguish between neighboring astrocyte cells. Subsequently, Figure 2(d) showcases the outcome of employing the Stardist plugin. While this algorithm excels in segmenting circular cells, it proves inadequate when confronted with more intricate cellular shapes, often failing to encompass the entire cell as a coherent entity. Finally, in Figure 2(e), an analogous behavior to the classic watershed is observed, with the method falling short in identifying cell boundaries, resulting in the inaccurate segmentation of neighboring cells.

**Fig 2.**
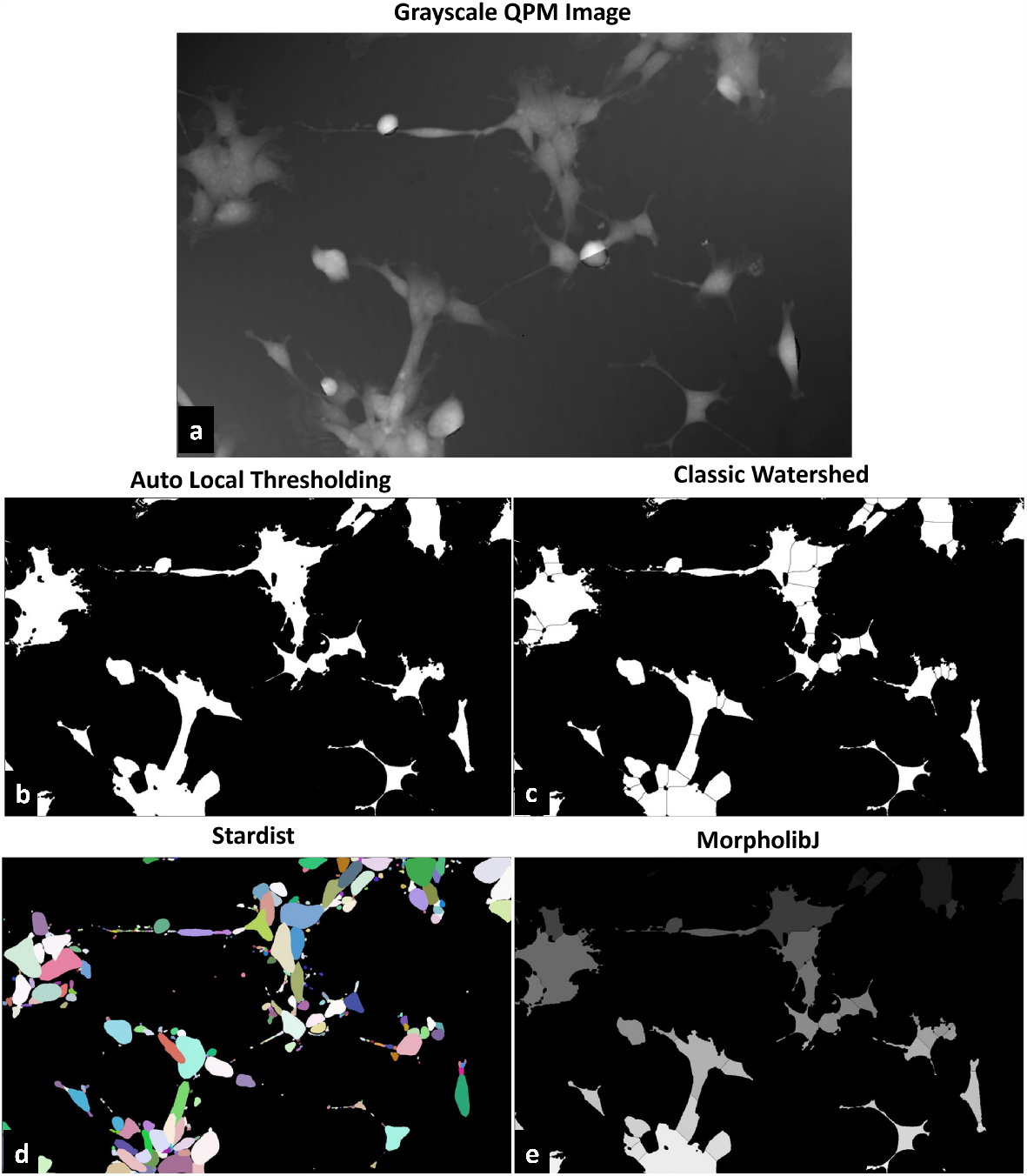
Comparison of various cell segmentation approaches on a representative FOV. a) The original grayscale FOV. b) Application of auto local thresholding and subsequent post-processing to eliminate small particles. c) Output of the classic watershed method. d) Segmentation achieved with the Stardist approach. e) Result obtained using the MorpholibJ algorithm.

Table 1 provides a comprehensive comparison of the performance metrics for Classic Watershed, MorphoLibJ, and Stardist in terms of correctly segmented cell count and neighboring cell count.

**Table 1.** Comparison of Correctly Segmented Cell % and Neighboring Cell % for Classic Watershed, MorphoLibJ, and Stardist

| Method | Correctly Segmented Cell % | Neighboring Cell % |
| --- | --- | --- |
| Classic Watershed | 12% | 11% |
| MorphoLibJ | 12% | 22% |
| Stardist | 6% | 55% |

In the realm of correctly segmented cell count, the evaluation was centered around the precision of boundary detection, emphasizing the accurate segmentation of cells within each image and processes recognition. Classic Watershed and MorphoLibJ demonstrated comparable results, exhibiting a higher count of correctly segmented cells compared to Stardist. However, all three of the programs struggled with catching the majority of the specific processes of the astrocytes, resulting in a low cell count percentage.

The other criteria tested was the neighboring cell count category, where the assessment focused on the precision of segmenting cells within specific astrocyte clusters. Stardist outperformed Classic Watershed and MorphoLibJ by more than twofold in effectively delineating individual astrocytes within clusters of cells. These results suggest that Stardist excels at discerning intricate structures and nuances within closely associated cell groups.

The findings underscore the strengths and weaknesses of each method, with Classic Watershed and MorphoLibJ excelling in overall cell segmentation precision, while Stardist stands out for its proficiency in segmenting individual astrocytes within clusters.

### 1.2 Final cell segmentation procedure

After processing the interference cell images for each cell culture dish at various time points into final real height maps, they are loaded as image sequences in ImageJ. At each time point, rectangles are drawn manually around the cells to determine their bounding box. The ‘measure’ tool located in the ‘analyze’ section is used to measure the position, height, and width of each rectangle. The resulting data is saved as CSV files and loaded in MATLAB to crop the real height map images. The cropped images, each containing one cell at a certain time point are then batch-processed using the following code in ImageJ, to create masks: [language=Java, basicstyle=] run(“Auto Local Threshold”, “method=Median radius=50 parameter_1_= 0*parameter*_2_= 0*white*”); *setOption*(“*BlackBackground*”, *true*); *run*(“*Erode*”); *run*(“*Dilate*”); This code applies the “Auto Local Threshold” on images, using the “Median” method with a 50-pixel radius. The process is followed by one round of Erosion to eliminate noise and small particles, and one round of Dilation. When cells within the bounding box are closely packed, the process of isolating the desired cell may lead to generating more than one object in the mask. One solution is to import the masks into MATLAB for further image processing. The ‘regionprops’ function is used to identify the object with the highest surface area, which corresponds to the main cell in the image. The other objects are eliminated, resulting in a mask with only one object that corresponds to the desired cell. Figure 3 displays two cells and their respective masks. Each cell has three corresponding images: a grayscale image, a mask image created with ImageJ, and a final processed image generated with MATLAB.

**Fig 3.**
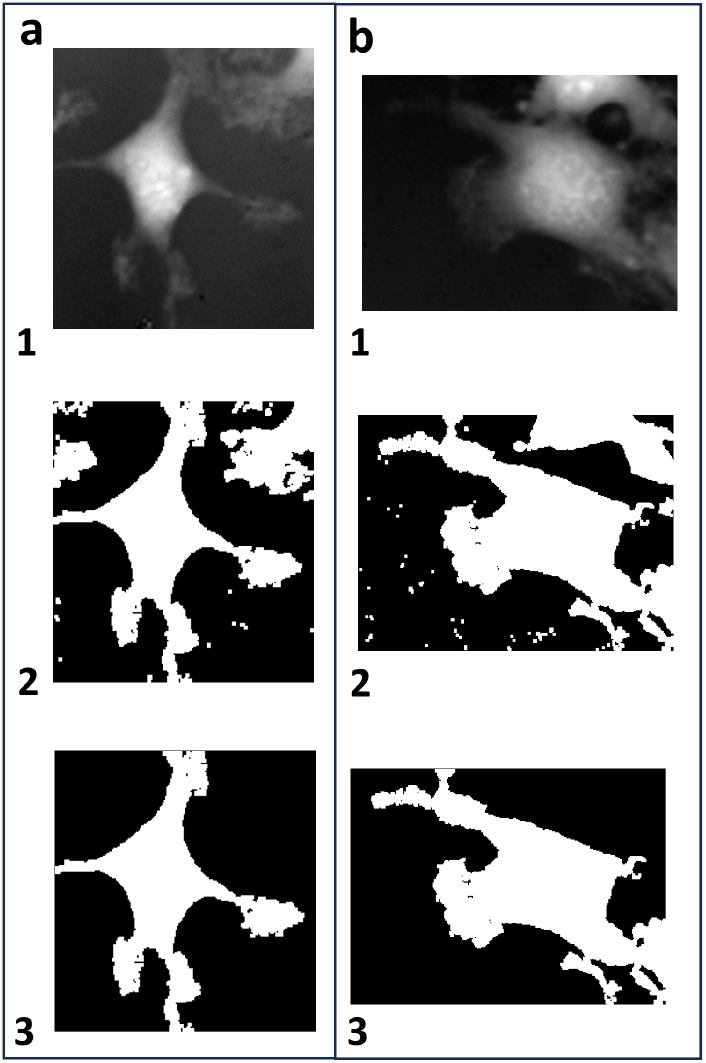
Two example cells (a,b) and their respective masks. Each cell has three corresponding images: (1) a grayscale image, (2) a mask image created with ImageJ, and (3) a final processed image generated with MATLAB.

### 1.3 Background correction

To eliminate any background interference caused by uneven exposure, a mask generated through ImageJ processing is utilized as a guide for calculating the average background height. The mask includes all particles within the bounding box, and the remaining part of the image represents the background. The calculated average is then subtracted from the entire image, resulting in a background-corrected image shown in Figure 4. Figures 4(a,b) display the resulting masks obtained from ImageJ and processed using MATLAB, respectively. Figures 4(c,d) show the original cell image and the background-corrected image, respectively, along with the height value of a point on the background provided for reference in both images.

**Fig 4.**
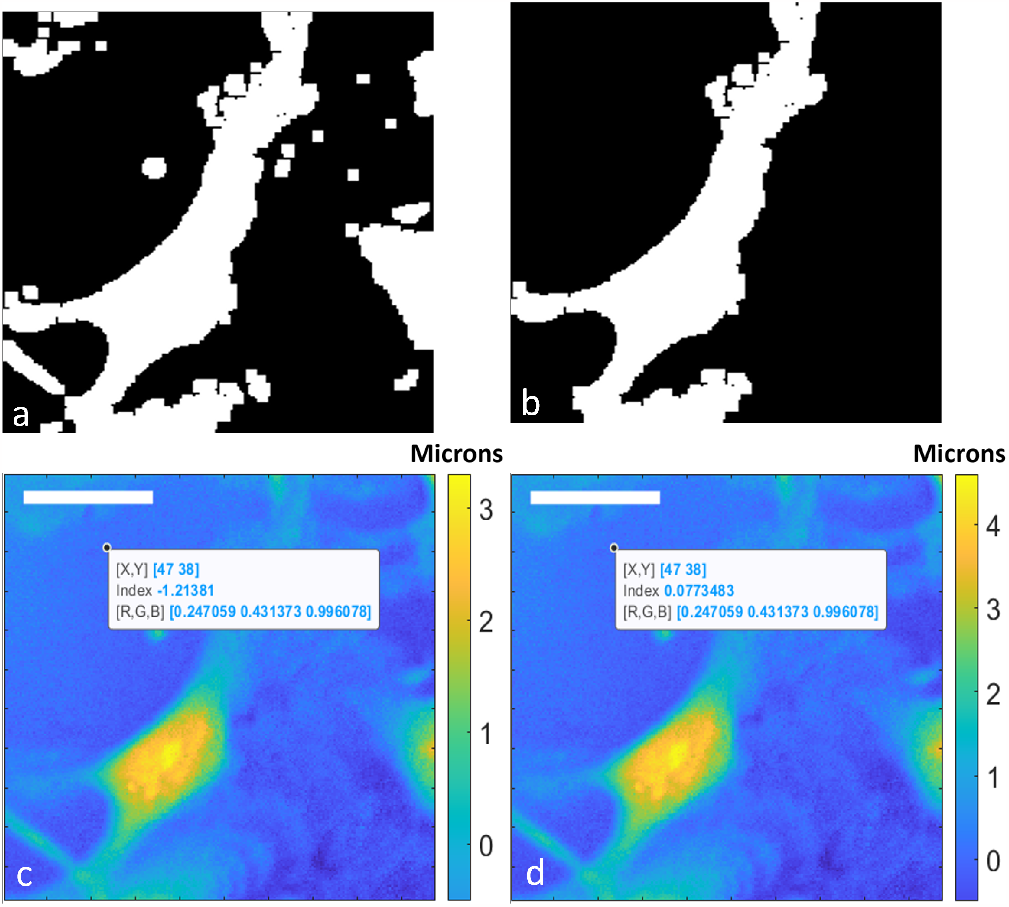
(a) The resulting masks obtained from ImageJ, (b) The mask processed using MATLAB, (c) the original cell image d) the background-corrected image. The height value of a point on the background is provided for reference in both images. The scale bar is 20 microns.

### 1.4 Feature extraction

Several morphological features are measured from the phase and height map images, including dry mass (DM), dry mass surface density (DMSD), surface area (SA), surface area to volume ratio (SAV), surface area to dry mass ratio (SADM), sphericity index, Height variance (HV), Height kurtosis (HK), Height skewness (HS), eccentricity (Ecc), volume divided by area (VoldivArea) and circularity (Cir). All features, except volume divided by area and circularity, are based on the parameters described in [27]. A detailed explanation of the features is described below:

- **Dry Mass (DM)**: The dry mass of the cell is proportional to its height profile. The relationship is determined by the refractive increment (*α*), which is typically constant for the type of cell under investigation. This measurement is crucial in biological research as it helps scientists understand the cellular composition and relative proportions of different components. Dry mass is calculated using the formula:

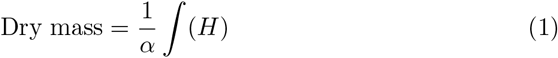

where *H* is the height of the cell at each point. For astrocytes, *α* can be estimated as 1.8 *×* 10^−4^ m^3^ kg^−1^.
- **Dry Mass Surface Density (DMSD)**: This is the dry mass per unit area. It’s calculated as:

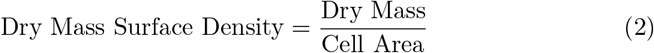
- **Volume**: The volume is calculated by integrating the height profile over the cell area:

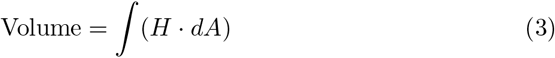

where *dA* is the differential area element. The volume of a cell provides insights into its overall size and shape. It is a fundamental parameter for understanding cell biology.
- **Surface Area from Height Profile (SA)**: This is derived from the height profile and provides information about the variation in the cell’s surface.
- **Projected Area (PA)**: The cell’s projected area is calculated by counting the number of pixels within the cell mask and multiplying it by the pixel size. This provides the two-dimensional surface area. PA is used to assess the cell’s planar size.
- **Surface Area to Volume Ratio (SAV)**: This ratio is obtained by dividing the surface area by the volume calculated earlier. The SAV ratio offers insight into the relationship between the surface area of a cell and its internal content, as a higher SAV may indicate a more extensive cell membrane relative to its volume.
- **Surface Area to Dry Mass Ratio**: Similar to the above, this ratio is found by dividing the surface area by the dry mass and provides insight into the relationship between the surface area of a cell and its internal content. The Surface Area to Dry Mass Ratio reflects the distribution of dry mass across the cell’s surface area.
- **Sphericity Index**: The sphericity index measures how closely the cell approximates a perfect sphere. It is calculated using the formula:

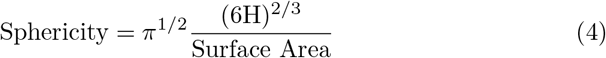

A perfect sphere has a sphericity value of 1. Sphericity is essential for understanding cell shape and deviations from a spherical structure, as cells with a higher sphericity index tend to be more spherical in shape.
- **Height Variance (HV)**: Height Variance (HV) is a parameter that measures the spread of height values within the cell. It provides insights into how varied the heights of different regions within the cell are. HV can be calculated as the variance of the height values over the cell area using the following formula:

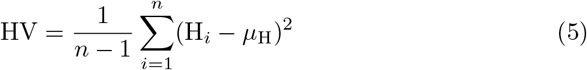

Where: - HV is the Height Variance. - *n* is the number of data points (height values). - H_*i*_ represents the individual height value. - *μ*_H_ is the mean of the height values. Higher HV values indicate greater height variation within the cell, potentially reflecting structural complexities or cellular processes.
- **Height Kurtosis (HK)**: Height Kurtosis (HK) is a parameter that measures whether the distribution of heights within the cell (height distribution) is more peaked or flat compared to a normal distribution. It quantifies the shape of the height distribution. HK can be calculated using the formula for kurtosis:

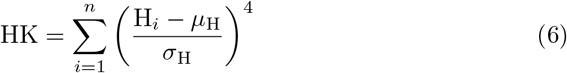

Where: - HK is the Height Kurtosis. - *n* is the number of data points (height values). - H_*i*_ represents the individual height value. - *μ*_H_ is the mean of the height values. - *σ*_H_ is the standard deviation of the height values. Deviations from a normal distribution can indicate irregularities in cell structure.
- **Height Skewness (HS)**: Height Skewness (HS) is a parameter that quantifies the asymmetry of the height distribution within the cell. It describes whether the distribution is skewed to the left or right. HS can be calculated using the formula for skewness:

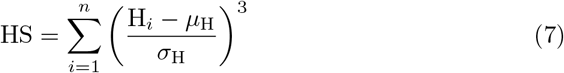

Where: - *n* is the number of data points (height values). - H_*i*_ represents the individual height value. - *μ*_H_ is the mean of the height values. - *σ*_H_ is the standard deviation of the height values. Changes in HS may suggest structural alterations.
- **Cell Eccentricity**: Eccentricity quantifies how much the cell’s shape deviates from a perfect circle. It is determined by finding the maximum and minimum radii (*r*_max_ and *r*_min_) of the cell’s boundary and applying the formula:

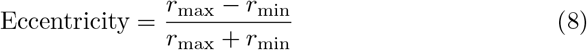

Deviations from a circular shape can provide information about cellular processes or structural changes.
- **Projected Area to Volume Ratio (PAV)**: This ratio quantifies cell flatness. It is calculated by dividing the projected area (2D area) by the volume of the cell (3D measurement). This allows researchers to assess whether cells are more planar or three-dimensional in shape.
- **Circularity**: Determined using the circularity feature in MATLAB’s ‘regionprops’ function. It evaluates the similarity of the cell’s shape to that of a perfect circle. This measure ranges from 0 to 1, with 1 representing a perfect circle. Higher circularity values indicate a closer resemblance to a circular shape. Deviations from circularity can suggest alterations in cell structure.
- **Perimeter (Perim)** Perimeter (perim) is a feature that represents the total length of the outer boundary of a cell. It provides information about the cell’s extent and boundary complexity. Changes in the perimeter may indicate variations in cell size or irregularities in cell shape.
- **Perimeter-to-Area Ratio (PerimDivArea)** PerimDivArea is a feature calculated by dividing the perimeter of a cell by its area. This ratio offers insight into the cell’s boundary irregularity relative to its overall size. Higher values may suggest a more complex and convoluted cell boundary compared to its size.
- **Complexity Score**: Computed as the ratio of the difference between the area of the cell’s convex hull and the cell area to the cell area itself. A lower score signifies a smoother and less intricate cell shape.

These features collectively contribute to our understanding of cell biology, enabling researchers to assess the effects of various conditions, treatments, or genetic manipulations on cellular structure and function.

Table 2 shows the corresponding feature values for each image along with a grayscale image of a cell at two different time points and their corresponding final masks, represented in Figure 5.

**Table 2.**
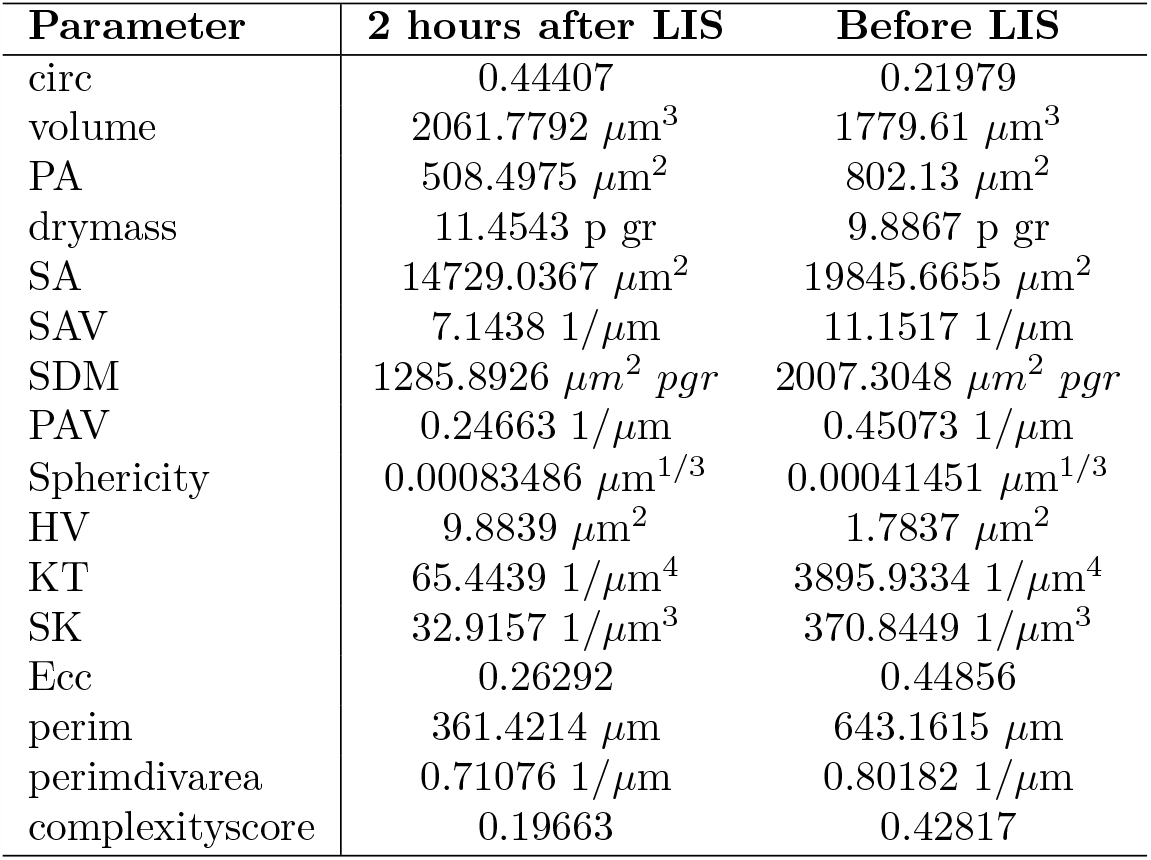
Comparison of Features for Cell Shown in Figure 5, Before and 2 Hours After LIS

| Parameter | 2 hours after LIS | Before LIS |
| --- | --- | --- |
| circ | 0.44407 | 0.21979 |
| volume | 2061.7792 $\mu\text{m}^3$ | 1779.61 $\mu\text{m}^3$ |
| PA | 508.4975 $\mu\text{m}^2$ | 802.13 $\mu\text{m}^2$ |
| drymass | 11.4543 p gr | 9.8867 p gr |
| SA | 14729.0367 $\mu\text{m}^2$ | 19845.6655 $\mu\text{m}^2$ |
| SAV | 7.1438 $1/\mu\text{m}$ | 11.1517 $1/\mu\text{m}$ |
| SDM | 1285.8926 $\mu\text{m}^2 \text{ pgr}$ | 2007.3048 $\mu\text{m}^2 \text{ pgr}$ |
| PAV | 0.24663 $1/\mu\text{m}$ | 0.45073 $1/\mu\text{m}$ |
| Sphericity | 0.00083486 $\mu\text{m}^{1/3}$ | 0.00041451 $\mu\text{m}^{1/3}$ |
| HV | 9.8839 $\mu\text{m}^2$ | 1.7837 $\mu\text{m}^2$ |
| KT | 65.4439 $1/\mu\text{m}^4$ | 3895.9334 $1/\mu\text{m}^4$ |
| SK | 32.9157 $1/\mu\text{m}^3$ | 370.8449 $1/\mu\text{m}^3$ |
| Ecc | 0.26292 | 0.44856 |
| perim | 361.4214 $\mu\text{m}$ | 643.1615 $\mu\text{m}$ |
| perimdivarea | 0.71076 $1/\mu\text{m}$ | 0.80182 $1/\mu\text{m}$ |
| complexityscore | 0.19663 | 0.42817 |

**Fig 5.**
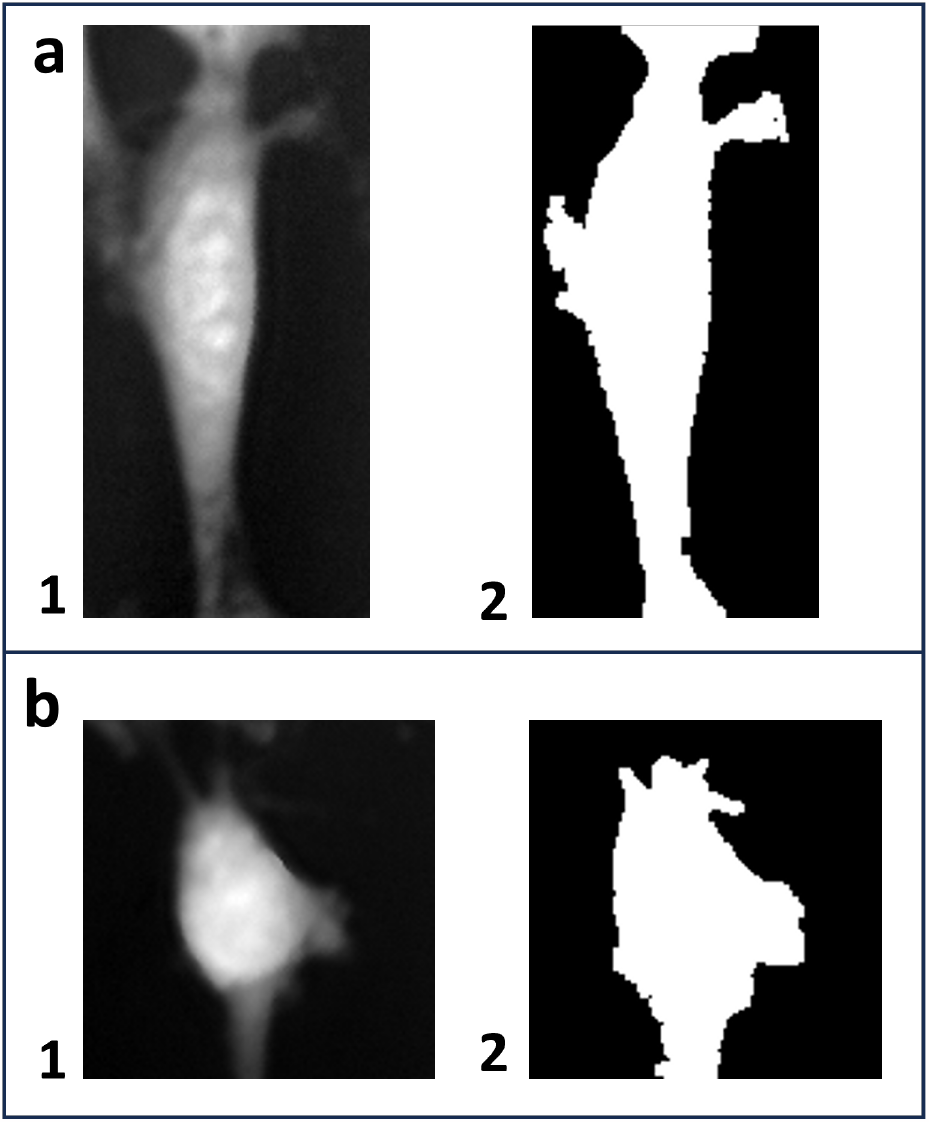
(a,b) Two grayscale images of a cell in two different time points: before LIS and 2 hours after LIS. (1,2) The corresponding final masks.

## 2 Experiments with the QPM’s current setup

### 2.1 Experiments protocol

#### 2.1.1 AST Cell Preparation

An established astrocyte type-I (Ast-1) line (clone CRL-2541) was received directly from ATCC. Ast-1 cells were cultured in advanced DMEM media supplemented with 2% FBS, 1% glutamax, and 0.2% gentamicin/amphotericin B. Ast-1 cells were plated onto glass-bottom 35mm imaging dishes. Alternatively, imaging dishes were coated by incubating them for 1 hour in a 1:60 dilution of matrigel:DMEM to promote cell adhesion.

#### 2.1.2 Control group

Astrocyte cells were maintained and observed within a temperature-controlled chamber designed specifically for the Quantitative Phase Microscope setup. The chamber’s environmental conditions, including the composition of the air, were precisely controlled using an Ibidi gas mixer, which ensured a stable and controlled atmosphere. The key parameters that were regulated included the humidity level, which was maintained at a constant 80 percent, and the carbon dioxide (CO2) concentration, which was set at 5%. These controlled conditions were essential to replicate the natural environment of these cells and minimize possible stress or alterations in their behavior.

During the observation period, the astrocyte cells were subjected to a time-lapse imaging protocol. Every 20 minutes, the cells were imaged and this process continued for a total duration of two hours. We ensured that there was no interference or disruption to the cells’ normal activity or behavior. This stringent approach allowed us to capture and analyze the astrocyte cells’ activities and interactions in their near-native state, providing valuable insights into their functions and responses under controlled conditions. In total, 84 cells from 4 separate dishes underwent analysis and subsequent processing for cell segmentation and feature extraction.

#### 2.1.3 Laser-induced shockwave group

The astrocyte cells were kept under similar conditions as the control group, except for being exposed to a laser-induced shockwave. This experiment involved a continuous imaging process, which started 20 minutes before the shockwave and lasted for 2 hours afterwards. The resulting quantitative phase images were analyzed to detect and measure various aspects of cell morphology.

To evaluate the effect of the shockwave on the cells, we focused on their morphology before and 10 minutes, and 2 hours after the shockwave. By doing so, we could identify any rapid or gradual changes or responses induced by the shockwave. This approach allowed us to gain a deeper understanding of how cells react and adapt to sudden environmental changes under these unique experimental conditions. A total of 204 cells collected from four dishes were subjected to analysis.

### 2.2 Image processing

For cell selection, both the cells that survived the shockwave and the cells fully within the field of view (FOV) were manually chosen using ImageJ. The detection of black and white masks for each individual cell was achieved by applying local thresholding with the median method in ImageJ. These masks were then imported into MATLAB for further image processing. The remaining image processing tasks were performed using MATLAB.

The cells were categorized based on their distance from the center of the shockwave, specifically into two groups: those located within 300 microns and those located beyond 300 microns.

Different morphological characteristics were examined for each group, including flatness, roundness, volume, volume divided by area, and circularity. The volume of a cell is determined by summing the thickness values. This measure provides information about the overall size and three-dimensional structure of the cell, which can be valuable in understanding cellular morphology and changes resulting from the shockwave impact. The area is calculated by counting the number of pixels within the cell mask and multiplying it by the pixel size. This provides a quantitative measurement of the cell’s two-dimensional surface area. It helps assess the cell’s coverage and can be used in conjunction with other features to analyze cell shape and compactness. Roundness is defined as 4 times pi multiplied by the ratio of the volume to the square of the surface area. This measure describes the degree to which the cell approximates a perfect sphere. It provides insights into the cell’s overall shape and can be useful in characterizing changes in cell morphology induced by the shockwave. Flatness is calculated by dividing the maximum Feret diameter (the longest distance between any two points on the cell’s boundary) by the minimum Feret diameter (the shortest distance between any two points on the cell’s boundary). This measurement quantifies the degree of elongation or flattening of the cell. It helps assess the cell’s shape and can indicate deformations caused by the shockwave. Circularity is determined using the circularity feature in MATLAB’s regionprops function. It evaluates the similarity of the cell’s shape to that of a perfect circle. This measure ranges from 0 to 1, with 1 representing a perfect circle. Circular cells will have higher circularity values, while irregularly shaped cells will have lower values. It helps assess the cell’s symmetry and can indicate any distortion or alteration resulting from the shockwave impact.

## 3 Statistical analysis

The statistical tests conducted to assess the changes in astrocyte cell parameters within the control and LIS group were performed through MATLAB, using the Wilcoxon signed-rank test, a non-parametric test suitable for paired data. Since the measurements were obtained from the same set of cells during the imaging period, the data are inherently dependent on each other. The Wilcoxon signed-rank test is well-suited for such paired comparisons, providing a robust analysis that does not assume a normal distribution of the data. This method accounts for the paired nature of the observations, making it appropriate for detecting subtle changes in the astrocyte cell parameters over time.

The Mann-Kendall Tau-b test [28] was also performed for data trend analysis. This is a non-parametric statistical method employed for the detection of monotonic trends in time series data. It specifically assesses the presence of an upward or downward trend over time without assuming any underlying distribution of the data. The Taub statistic represents the magnitude of the trend, and the p-value indicates the statistical significance of that trend.

### 3.1 Control group

In the control group, astrocyte cells were subjected to two hours of imaging, and the average value for each feature for the total of 125 cells, is summarized in Table 3.

**Table 3.** AST1 Cells Various Features Average Values

| Parameters | Average Value (n=84) |
| --- | --- |
| Circularity | 0.24813 |
| Volume | 3054.7392 $\mu\text{m}^3$ |
| Projected Area | 1125.4119 $\mu\text{m}^2$ |
| Dry Mass | 16.9708 p gr |
| Surface Area | 29107.4126 $\mu\text{m}^2$ |
| Surface Area to Volume Ratio | 9.6821 $1/\mu\text{m}$ |
| Surface Area to Dry Mass Ratio | 1742.7744 p gr/$\mu\text{m}^2$ |
| Projected Area to Volume Ratio | 0.37841 $1/\mu\text{m}$ |
| Sphericity | 0.00024686 $\mu\text{m}^{1/3}$ |
| Height Variance | 4.4972 $\mu\text{m}^2$ |
| Height Kurtosis | 4375.1226 $1/\mu\text{m}^4$ |
| Height Skewness | 879.4492 $1/\mu\text{m}^3$ |
| Eccentricity | 0.28863 |
| Perimeter | 833.5168 $\mu\text{m}$ |
| Perimeter to Area Ratio | 0.74369 $1/\mu\text{m}$ |
| Complexity Score | 0.38273 |

Various parameters were measured to assess any potential changes during this imaging period.

After two hours of imaging, a comparison of feature values was conducted, as presented in Table 4. The test statistics, p-values, and the determination of significant differences were reported for each feature. Despite this analysis, no statistically significant changes were observed in any of the features within the control group. The lack of statistical significance suggests that the astrocyte cells’ morphology and characteristics remained stable during the experimental timeframe.

**Table 4.** Significance Test Results for the Control Group in 2 Hours

| Parameters | P-Value | Sig. | Mean Diff. |
| --- | --- | --- | --- |
| Circularity | 0.71459 | No | 0.0066564 |
| Volume | 0.87245 | No | -5.8727 $\mu\text{m}^3$ |
| Projected Area | 0.71959 | No | -1.9396 $\mu\text{m}^2$ |
| Dry Mass | 0.87245 | No | -0.032626 p gr |
| Surface Area | 0.92538 | No | 316.3684 $\mu\text{m}^2$ |
| Surface Area to Volume Ratio | 0.56207 | No | 0.49498 $1/\mu\text{m}$ |
| Surface Area to Dry Mass Ratio | 0.56207 | No | 89.0972 $\mu\text{m}^2/\text{p gr}$ |
| Projected Area to Volume Ratio | 0.78559 | No | 0.017124 $1/\mu\text{m}$ |
| Sphericity | 0.96087 | No | 3.8919e-07 $\mu\text{m}^{1/3}$ |
| Height Variance | 0.057451 | No | 0.4453 $\mu\text{m}^2$ |
| Height Kurtosis | 0.43512 | No | 572.6861 $1/\mu\text{m}^4$ |
| Height Skewness | 0.9183 | No | -164.8099 $1/\mu\text{m}^3$ |
| Eccentricity | 0.2167 | No | -0.015709 |
| Perimeter | 0.99644 | No | -10.0189 $\mu\text{m}$ |
| Perimeter to Area Ratio | 0.23904 | No | -0.01669 $1/\mu\text{m}$ |
| Complexity Score | 0.10076 | No | -0.017269 |

In conclusion, the data from the control group indicate that, no significant changes occurred in the astrocyte cells after two hours of imaging. This stability in cellular features provides a baseline for comparison with experimental groups, allowing for a more accurate assessment of any observed effects or changes due to LIS.

**Table 5.** Statistical Parameters for the cells 1 Min Before LIS Vs. 10 Min After LIS

| Parameters | P-Value | Sig. | Mean Diff. |
| --- | --- | --- | --- |
| Circularity | 0.00056679 | Yes | 0.029897 |
| Volume | 1.51E-08 | Yes | -352.0049 $\mu\text{m}^3$ |
| Projected Area | 1.27E-10 | Yes | -167.5746 $\mu\text{m}^2$ |
| Dry Mass | 1.51E-08 | Yes | 1.9556 p gr |
| Surface Area | 2.62E-10 | Yes | -3708.6028 $\mu\text{m}^2$ |
| Surface Area to Volume Ratio | 9.60E-17 | Yes | -2.0859 $1/\mu\text{m}$ |
| Surface Area to Dry Mass Ratio | 9.60E-17 | Yes | -375.4581 $\mu\text{m}^2/\text{p gr}$ |
| Projected Area to Volume Ratio | 2.20E-29 | Yes | -0.088707 $1/\mu\text{m}$ |
| Sphericity | 4.83E-23 | Yes | 0.00019486 $\mu\text{m}^{1/3}$ |
| Height Variance | 8.99E-26 | Yes | 2.1892 $\mu\text{m}^2$ |
| Height Kurtosis | 3.47E-23 | Yes | -3588.9706 $1/\mu\text{m}^4$ |
| Height Skewness | 5.91E-07 | Yes | -580.7853 $1/\mu\text{m}^3$ |
| Eccentricity | 0.050085 | No | 0.0094032 |
| Perimeter | 3.87E-07 | Yes | -128.5676 $\mu\text{m}$ |
| Perimeter to Area Ratio | 0.23904 | No | -0.01669 $1/\mu\text{m}$ |
| Complexity Score | 0.10076 | No | -0.014833 |

## 4 Laser-Induced Shockwaves (LIS) group

The investigation of astrocyte responses to laser-induced shockwaves (LIS) has revealed profound morphological alterations, providing intricate insights into the dynamic cellular adaptations. Analyzing various morphological features immediately after LIS and 2 hours post-LIS has elucidated the immediate responses and the persistence of changes, offering a comprehensive understanding of astrocytic behavior under mechanical stress. The morphological changes observed in astrocytes following LIS reveal immediate adaptive responses and persistent alterations. The cells exhibit a rapid transition to a more circular shape, as evidenced by increased circularity. Simultaneously, an increase in volume and changes in surface characteristics, including surface area (SA) and surface area to volume ratio (SAV), signify dynamic adaptations aimed at withstanding the mechanical stress induced by LIS.

Height characteristics, such as height variance and non-uniformity, suggest rapid cytoskeletal rearrangements and structural adaptations. Furthermore, alterations in sphericity indicate a shift towards a more spherical cell shape, potentially enhancing cell stability.

A 2-hour post-LIS analysis reveals sustained changes in surface area related parameters, volume, and height characteristics, indicating ongoing cellular adjustments. The persistence of these alterations suggests a prolonged cellular response, possibly involving regulatory processes aimed at restoring homeostasis Table 6.

**Table 6.** Statistical parameters for the cells 10 min after LIS Vs. 2 Hr after LIS

| Parameters | P-Value | Sig. | Mean Diff |
| --- | --- | --- | --- |
| Circularity | 0.41917 | No | 0.010624 |
| Volume | 5.80E-12 | Yes | -398.7118 $\mu\text{m}^3$ |
| Projected Area | 6.72E-05 | Yes | 114.8401 $\mu\text{m}^2$ |
| Dry Mass | 5.80E-12 | Yes | -2.2151 p gr |
| Surface Area | 6.97E-07 | Yes | 2445.7871 $\mu\text{m}^2$ |
| Surface Area to Volume Ratio | 5.61E-21 | Yes | 2.8767 $1/\mu\text{m}$ |
| Surface Area to Dry Mass Ratio | 5.61E-21 | Yes | 517.8092 $\mu\text{m}^2/\text{p gr}$ |
| Area to Volume Ratio | 1.52E-25 | Yes | 0.092696 $1/\mu\text{m}$ |
| Sphericity | 2.35E-21 | Yes | -0.00016332 $\mu\text{m}^{1/3}$ |
| Height Variance | 2.18E-17 | Yes | -1.9532 $\mu\text{m}^2$ |
| Height Kurtosis | 8.76E-18 | Yes | 7017.1477 $1/\mu\text{m}^4$ |
| Height Skewness | 0.001124 | Yes | 538.0122 $1/\mu\text{m}^3$ |
| Eccentricity | 0.035607 | Yes | -0.015711 |
| Perimeter | 0.074821 | No | 47.807 $\mu\text{m}$ |
| Perimeter to Area Ratio | 5.31E-05 | Yes | -0.035515 $1/\mu\text{m}$ |
| Complexity Score | 0.0015005 | Yes | -0.032056 |

In Table 7, the statistical parameters provide a detailed insight into the morphological changes in astrocytes 1 minute before Laser-Induced Shockwave (LIS) compared to 2 hours after LIS. Notably, several features exhibit significant alterations during this time frame, shedding light on the dynamic responses of astrocytes to the mechanical perturbation induced by the shockwave.

**Table 7.** Statistical Parameters for the cells 1 Min Before LIS Vs. 2 Hr After LIS

| Parameters | P-Value | Sig. | Mean Diff |
| --- | --- | --- | --- |
| Circularity | 0.0001094 | Yes | 0.040521 |
| Volume | 0.034178 | Yes | -46.707 $\mu\text{m}^3$ |
| Projected Area | 0.082363 | No | -52.7344 $\mu\text{m}^2$ |
| Dry Mass | 0.034178 | Yes | -0.25948 p gr |
| Surface Area | 0.0026958 | Yes | -1262.8157 $\mu\text{m}^2$ |
| Surface Area to Volume Ratio | 0.21533 | No | 0.79084 $1/\mu\text{m}$ |
| Surface Area to Dry Mass Ratio | 0.21533 | No | 142.3511 $\mu\text{m}^2/\text{p gr}$ |
| Area to Volume Ratio | 0.33316 | No | 0.0039881 $1/\mu\text{m}$ |
| Sphericity | 0.024559 | Yes | 3.1541e-05 $\mu\text{m}^{1/3}$ |
| Height Variance | 0.057276 | No | 0.23605 $\mu\text{m}^2$ |
| Height Kurtosis | 0.011555 | Yes | 3428.1771 $1/\mu\text{m}^4$ |
| Height Skewness | 0.85525 | No | -42.7731 $1/\mu\text{m}^3$ |
| Eccentricity | 0.59318 | No | -0.0063078 |
| Perimeter | 0.001608 | Yes | -80.7606 $\mu\text{m}$ |
| Perimeter to Area Ratio | 0.0019983 | Yes | -0.028057 $1/\mu\text{m}$ |
| Complexity Score | 1.86E-06 | Yes | -0.046889 |

Among the noteworthy changes, the circularity of astrocytes increases significantly (*p*-value = 0.0001094), indicating a shift toward a more rounded cell shape.

Concurrently, there is a considerable decrease in both cell volume (*p*-value = 0.034178) and dry mass (*p*-value = 0.034178), suggesting potential cellular contraction or changes in density. The increase in sphericity (*p*-value = 0.024559) further supports a trend toward a more spherical cellular morphology, indicative of a response to mechanical stress or injury.

Examining the outer characteristics of the astrocytes, both perimeter (*p*-value = 0.001608) and perimeter-to-area ratio (*p*-value = 0.0019983) significantly decrease. This reduction implies a more compact cell structure or altered cell-cell interactions.

Additionally, the increase in kurtosis (*p*-value = 0.011555) suggests changes in the height distribution within the cell, indicating potential alterations in cellular structure or response to the shockwave.

The observed morphological adaptations may be integral to the cellular response mechanism, influencing cellular mechanics, cell-cell interactions, and potentially contributing to the overall functionality of astrocytes. Understanding these changes is crucial for deciphering the nuanced ways in which astrocytes respond to mechanical stress, and it opens avenues for further investigations into the underlying cellular processes driving these morphological alterations. The Mann-Kendall Tau-b test results (Table 9) underscore the presence of significant trends in various morphological features, consolidating the robustness of the observed changes. Key variables, such as volume and surface area features, and height variance, manifest noteworthy trends, emphasizing the biological significance of these alterations. Figure 6 represents the average values of surface area, surface area to volume ratio, volume, and projected area to volume ratio at different time points, measured in minutes after LIS. The fitted line in the figure was obtained using the fitlm function in MATLAB to fit a linear regression model. Upon analyzing Figure 6 and Table 9, we can observe a significant increase in surface area and surface area divided by volume. This peculiar trend may indicate a cellular response aimed at enhancing chemical interactions with the surrounding environment.

**Fig 6.**
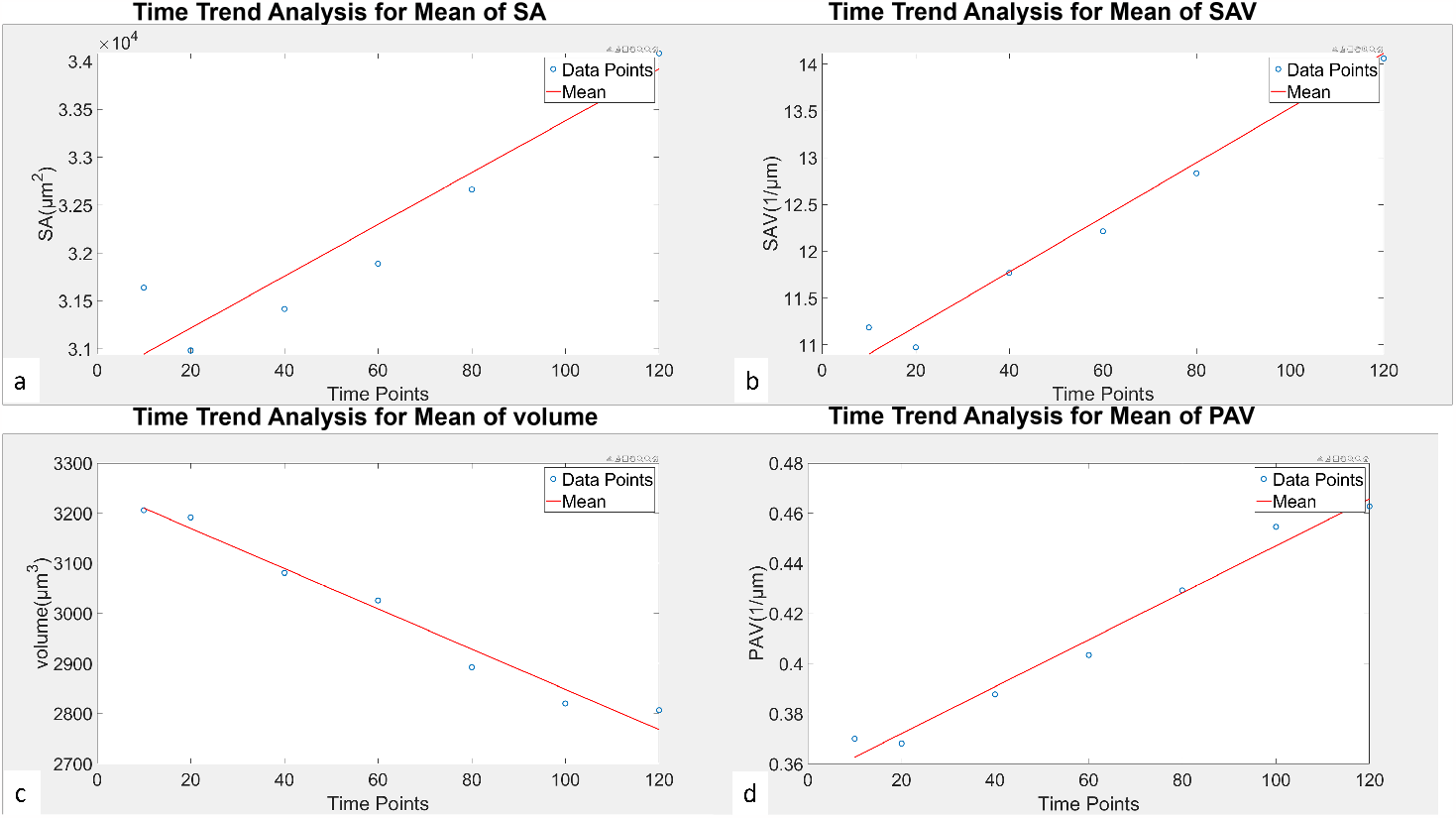
Time Trend Analysis for the mean value of various features. The time points show the time after LIS in minutes.

Additionally, it is noticeable that the projected area and projected area to volume ratio are also showing a significant increase over time, suggesting that the cells are getting flatter in response to LIS. These dynamics highlight the complexity of the cellular response to LIS, emphasizing the need for further investigation into the underlying biological mechanisms driving these morphological changes.

The observed morphological changes in astrocytes in response to LIS hold profound biological significance. The shift towards a more circular cell shape, reduced volume, and alterations in surface characteristics may indicate an orchestrated effort to enhance cell stability and mitigate the impact of mechanical stress. The simultaneous changes in height characteristics, such as increased variance and non-uniformity, suggest dynamic cytoskeletal rearrangements and structural adaptations to the shockwave.

Furthermore, the persistence of these morphological adaptations over time underscores the resilience and regulatory capacity of astrocytes. The maintained alterations in sphericity, eccentricity, and boundary features may reflect the ongoing role of astrocytes in responding to and mitigating mechanical perturbations, emphasizing their dynamic nature and ability to adapt to external stimuli.

Table 8 provides insights into the significant changes in cellular morphological features at various time points following LIS, compared to the state 1 minute before LIS. The highlighted cells (in red) indicate significant alterations in the respective features. The consistent significance in circularity and sphericity throughout the observed time points suggests a sustained change in cell circularity, indicating potential alterations in cell shape stability and dynamics. Features associated with the ratio of area to volume of the cells like SAV and PAV, or height distributions, such as variance, kurtosis, and skewness, exhibit significant changes post-LIS. However, after the 100 to 120-minute mark, no further significant changes are observed when compared to the pre-LIS measurements. This observation underscores the plasticity of the cells; despite undergoing substantial morphological changes and stress, they demonstrate recovery. However, the sustained significance in certain features underscores their potential role in cellular adaptation and functional implications during and after LIS-induced stress.

**Table 8.** Significance table of changes for each feature at various time points (in minutes) compared to 1 min before LIS

| Parameters/Time after LIS | 1" | 10" | 20" | 40" | 60" | 80" | 100" | 120" |
| --- | --- | --- | --- | --- | --- | --- | --- | --- |
| Circularity | 1 | 1 | 1 | 1 | 1 | 1 | 1 | 1 |
| Volume | 0 | 1 | 1 | 1 | 0 | 0 | 1 | 1 |
| Projected Area | 1 | 1 | 1 | 1 | 1 | 1 | 1 | 0 |
| Dry Mass | 0 | 1 | 1 | 1 | 0 | 0 | 1 | 1 |
| Surface Area | 1 | 1 | 1 | 1 | 1 | 1 | 1 | 1 |
| Surface Area to Volume Ratio | 1 | 1 | 1 | 1 | 1 | 1 | 0 | 0 |
| Surface Area to Dry Mass Ratio | 1 | 1 | 1 | 1 | 1 | 1 | 0 | 0 |
| Area to Volume Ratio | 1 | 1 | 1 | 1 | 1 | 1 | 0 | 0 |
| Sphericity | 1 | 1 | 1 | 1 | 1 | 1 | 1 | 1 |
| Height Variance | 1 | 1 | 1 | 1 | 1 | 1 | 0 | 0 |
| Height Kurtosis | 1 | 1 | 1 | 1 | 1 | 0 | 0 | 1 |
| Height Skewness | 1 | 1 | 1 | 1 | 1 | 1 | 1 | 0 |
| Eccentricity | 0 | 0 | 0 | 0 | 0 | 0 | 0 | 0 |
| Perimeter | 1 | 1 | 1 | 1 | 1 | 1 | 1 | 1 |
| Perimeter to Area Ratio | 0 | 0 | 0 | 0 | 0 | 1 | 1 | 1 |
| Complexity Score | 0 | 0 | 0 | 1 | 1 | 1 | 1 | 1 |

**Table 9.** Mann-Kendall Tau-b Test Results. This test shows whether there is significant trend for a feature over the period of 20 minutes to 2 hours after LIS.

| Parameters | Tau-b | P-Value | Trend |
| --- | --- | --- | --- |
| Circularity | 0.3333 | 0.4524 | No significant trend detected |
| Volume | -1 | 0.0085 | Significant trend detected |
| Projected Area | 1 | 0.0085 | Significant trend detected |
| Dry Mass | -1 | 0.0085 | Significant trend detected |
| Surface Area | 1 | 0.0085 | Significant trend detected |
| Surface Area to Volume Ratio | 1 | 0.0085 | Significant trend detected |
| Surface Area to Dry Mass Ratio | 1 | 0.0085 | Significant trend detected |
| Area to Volume Ratio | 1 | 0.0085 | Significant trend detected |
| Sphericity | -1 | 0.0085 | Significant trend detected |
| Height Variance | -1 | 0.0085 | Significant trend detected |
| Height Kurtosis | 1 | 0.0085 | Significant trend detected |
| Height Skewness | 0.4667 | 0.2597 | No significant trend detected |
| Eccentricity | -0.4667 | 0.2597 | No significant trend detected |
| Perimeter | 0.6 | 0.1329 | No significant trend detected |
| Perimeter to Area Ratio | -0.7333 | 0.0603 | No significant trend detected |
| Complexity Score | -0.8667 | 0.0242 | Significant trend detected |

In summary, the expanded analysis of astrocyte responses to LIS reveals a multifaceted and dynamic cellular adaptation to mechanical stress. The initial acute responses are followed by sustained alterations, suggesting a complex interplay of protective mechanisms and cellular remodeling. The interconnectedness of morphological features, temporal dynamics, and functional implications underscore the intricate nature of astrocyte responses and pave the way for further exploration into the molecular mechanisms governing these adaptations. This expanded interpretation enhances our understanding of astrocyte biology in the context of mechanical perturbations and lays the groundwork for future investigations into the broader implications for central nervous system dynamics.

## 5 Discussion

The ability to extract features such as cell area, dry mass, and volume from quantitative phase images is crucial in cell biology [29]. These features offer quantitative and reproducible metrics for understanding various cellular characteristics, opening doors to a plethora of applications. The analysis of these features can serve as the foundation for comprehending cellular injuries, cell behavior, cell cycle progression, and responses to external stimuli or drugs.

These methods are highly relevant and useful in the study of various diseases like Traumatic Brain Injury (TBI), Huntington’s disease, and other neurodegenerative disorders. These methods help researchers extract precise cellular features, which in turn enable them to gain critical insights. For example, in the case of TBI, these techniques assist in assessing the extent of axonal injury, measuring the morphological changes in neurons and astrocytes, and keeping track of repair processes. Similarly, in the case of Huntington’s disease, which is characterized by protein aggregations within neurons, these methods help quantify changes in neuronal morphology caused by these aggregations. [29].

While the methods discussed in this chapter are highly effective for many aspects of cell biology research, there are challenges in the context of cell segmentation, particularly when dealing with astrocytes. Astrocytes play a vital role in the central nervous system and exhibit dynamic morphological changes in response to various stimuli [30].

Accurate quantification of astrocyte reactivity necessitates precise cell segmentation and the identification of features distinguishing reactive from non-reactive cells. Astrocyte cell segmentation presents two significant challenges. The first is creating a mask that distinctly separates cell borders, and the second is defining the borders of cell clusters. To address these challenges, we adjusted parameters, including thresholding parameters in ImageJ. While results were generally acceptable for most cells, there is room for improvement. For instance, in Figure 3, the final mask for cell ‘a’ accurately delineates cell borders, but for cell ‘b’, the neighboring cell on the bottom right remains unmasked. Additionally, the developed algorithm necessitates manual outlining of cell bounding boxes, which enhances accuracy but is time-consuming. A combination of thresholding the entire image using ImageJ, followed by a watershed algorithm and particle detection with ‘regionprops’ in MATLAB, was employed to automate bounding box outlining.

However, the watershed algorithm often led to poor border masking of astrocytes. Regarding feature extraction, our results show promise in extracting various features from cells. For example, comparing the cell before a shockwave (Figure 5(a)) with the cell two hours after the shockwave (Figure 5(b)), as presented in Table 2, reveals changes in eccentricity and circularity, indicative of elongation. However, astrocyte reactivity is more complex, requiring the development of features capable of capturing intricate morphological changes.

A potential future step involves the development of features that account for the quantity of astrocyte processes, such as fractal analysis. Fractal analysis quantifies the self-similarity and complexity of structures. Applying this technique to astrocyte cell morphology may offer valuable insights into their reactivity. The fractal dimension and other fractal parameters can measure the intricacy of astrocyte processes and detect reactivity by observing changes in these features over time. This approach transcends basic shape and size measurements, providing a more detailed and dynamic view of astrocyte behavior.

Deep learning and machine learning techniques have shown promise in addressing cell detection and feature extraction challenges [31, 32]. These methods have the advantage of being able to adapt to data patterns, reducing the need for complex handcrafted algorithms [31]. However, there are several hurdles associated with these techniques. One major challenge is the requirement for extensive and diverse training data. Deep learning models rely on large amounts of labeled data to learn and generalize effectively. Obtaining such datasets can be time-consuming and resource-intensive. Another challenge is the fine-tuning of model parameters and architecture. Finding the optimal configuration for a deep learning model can be a complex and iterative process. It often requires experimentation and computational resources to train and evaluate different models. Interpretability is also a concern when using deep learning techniques. These models often function as black boxes, making it difficult to understand the reasoning behind their decisions. This lack of interpretability can be problematic, especially in critical applications where explanations are necessary. Furthermore, deep learning methods are not one-size-fits-all and may require customization for specific tasks and datasets. Different cell detection and feature extraction tasks may have unique requirements and characteristics that need to be considered when designing and training deep learning models [31].

Despite these challenges, deep learning and machine learning techniques have shown promising results in various domains, including computational biology and medical imaging [31, 32]. With further advancements and experience, it is expected that these techniques will continue to evolve and become more accessible for addressing cell biology challenges.

This study aims to explore the morphological changes occurring in astrocyte cells in response to laser-induced shockwaves, providing valuable insights into the dynamics of cellular responses to mechanical stimuli. The motivation behind this research lies in comprehending the microscopic consequences of traumatic events, particularly traumatic brain injury (TBI), at the cellular level. Focusing on astrocytes, the most abundant cells in the central nervous system, we simulate TBI conditions in vitro by subjecting cells to controlled shear stress through laser-induced shockwaves. For accurate measurement of morphological alterations, we utilize quantitative phase microscopy (QPM). Our specific interest in astrocyte cells stems from their propensity to undergo astrogliosis, a triggered response to injury, with previous studies establishing morphological changes associated with this process. This research framework provides a coherent exploration of how astrocyte cells adapt morphologically to mechanical stress, shedding light on potential implications for understanding cellular responses to traumatic brain injuries. The experimental setup, involving a control group and a laser-induced shockwave group, ensures a systematic exploration of the effects of shockwaves on astrocyte morphology. For the control group, environmental conditions, such as humidity and CO2 levels, were controlled in order to minimize external factors that could influence cell behavior.

Image processing techniques, including phase unwrapping and cell segmentation, and feature extraction contribute to the precision of data analysis. Utilizing the QPM images of astrocyte cells comprehensive characteristics of astrocytes were measured, including an average height profile and average of parameters, such as dry mass, volume, and circularity, provide a comprehensive view of astrocyte morphology.

The experiments’ protocol, including the control and laser-induced shockwave groups, ensures a systematic and controlled approach to studying astrocyte responses. The use of time-lapse imaging and careful monitoring of environmental conditions enhances the reliability of the data. The results from the control group indicate stability in astrocyte morphology over the two-hour imaging period. This provides a crucial baseline for comparing with the laser-induced shockwave group, where dynamic and sustained changes are observed.

The investigation into astrocyte responses to laser-induced shockwaves (LIS) uncovers immediate and sustained morphological changes, providing insights into dynamic cellular adaptations. Following LIS, astrocytes rapidly transition to a more circular shape, accompanied by changes in volume, surface characteristics, and height features. A 2-hour post-LIS analysis reveals persistent alterations, suggesting ongoing cellular adjustments and regulatory processes. Statistical analysis highlights significant changes in circularity, volume, surface area, and other features, emphasizing dynamic responses to mechanical perturbation. Mann-Kendall Tau-b test results confirm the robustness of observed trends. Surface area dynamics suggest a cellular response to enhance chemical interactions, while increased projected area and area-to-volume ratio indicate flattening in response to LIS. Overall, these morphological changes signify astrocytes’ multifaceted and adaptive nature, enhancing our understanding of their responses to mechanical stress and paving the way for further exploration into molecular mechanisms and broader implications for the central nervous system. Biologically, these findings indicate a remarkable plasticity in astrocytes, actively adapting to mechanical stressors. The sustained changes in various features suggest intricate molecular and cellular mechanisms at play. Further investigations into mechanotransduction pathways are crucial to unravel the underlying processes driving these observed adaptations.

## Conclusion

In conclusion, the methods discussed in this paper empower researchers with powerful tools to extract valuable features from quantitative phase images, enhancing our understanding of cellular behavior. Moreover, for astrocyte cells and similar cases where reactivity plays a crucial role, exploring novel features can open new avenues for quantifying complex cellular responses. Laser-induced shockwaves induce dynamic and lasting morphological changes in astrocytes, shedding light on the intricate cellular responses to mechanical stimuli.

Overall, the scientific rigor, systematic approach, and thorough analysis presented contribute to the advancement of knowledge in the field of astrocyte biology and cellular responses to mechanical stimuli. The findings have implications for traumatic brain injuries and provide a foundation for future research in this critical area of neuroscience.

## Notes

### Competing Interest Statement

The authors have declared no competing interest.

## References

1. Ghajar J. Traumatic brain injury. The Lancet. 2000;356(9233):923–929.

2. Finan JD. Biomechanical simulation of traumatic brain injury in the rat. Clinical Biomechanics. 2019;64:114–121.

3. Nakagawa A, Fujimura M, Kato K, Okuyama H, Hashimoto T, Takayama K, et al. Shock wave-induced brain injury in rat: novel traumatic brain injury animal model. In: Acta Neurochirurgica Supplements. Springer; 2008. p. 421–424.

4. Selfridge A, Preece D, Gomez V, Shi L, Berns M. A model for traumatic brain injury using laser induced shockwaves. In: Optical Trapping and Optical Micromanipulation XII. vol. 9548. SPIE; 2015. p. 154–164.

5. Wakida NM, Cruz GMS, Ro CC, Moncada EG, Khatibzadeh N, Flanagan LA, et al. Phagocytic response of astrocytes to damaged neighboring cells. PloS one. 2018;13(4).

6. Burda JE, Bernstein AM, Sofroniew MV. Astrocyte roles in traumatic brain injury. Experimental neurology. 2016;275:305–315.

7. Bush TG, Puvanachandra N, Horner CH, Polito A, Ostenfeld T, Svendsen CN, et al. Leukocyte infiltration, neuronal degeneration, and neurite outgrowth after ablation of scar-forming, reactive astrocytes in adult transgenic mice. Neuron. 1999;23(2):297–308.

8. Codeluppi SA, Chatterjee D, Prevot TD, Bansal Y, Misquitta KA, Sibille E, et al. Chronic stress alters astrocyte morphology in mouse prefrontal cortex. International Journal of Neuropsychopharmacology. 2021;24(10):842–853.

9. Minge D, Domingos C, Unichenko P, Behringer C, Pauletti A, Anders S, et al. Heterogeneity and development of fine astrocyte morphology captured by diffraction-limited microscopy. Frontiers in cellular neuroscience. 2021;15:669280.

10. Bushong EA, Martone ME, Ellisman MH. Maturation of astrocyte morphology and the establishment of astrocyte domains during postnatal hippocampal development. International Journal of Developmental Neuroscience. 2004;22(2):73–86.

11. Phatnani H, Maniatis T. Astrocytes in neurodegenerative disease. Cold Spring Harbor perspectives in biology. 2015;7(6):a020628.

12. Anderson MA, Ao Y, Sofroniew MV. Heterogeneity of reactive astrocytes. Neuroscience letters. 2014;565:23–29.

13. Hu C, Popescu G. Quantitative phase imaging (QPI) in neuroscience. IEEE Journal of Selected Topics in Quantum Electronics. 2018;25(1):1–9.

14. Park Y, Depeursinge C, Popescu G. Quantitative phase imaging in biomedicine. Nature Photonics. 2018;12(10):578–589.

15. Yamauchi T, Iwai H, Yamashita Y. Label-free imaging of intracellular motility by low-coherent quantitative phase microscopy. Optics express. 2011;19(6):5536–5550.

16. Wakida NM, Cruz GMS, Pouladian P, Berns MW, Preece D. Fluid Shear Stress Enhances the Phagocytic Response of Astrocytes. Frontiers in Bioengineering and Biotechnology. 2020;8:1290.

17. Maneshi MM, Sachs F, Hua SZ. Heterogeneous cytoskeletal force distribution delineates the onset Ca2+ influx under fluid shear stress in astrocytes. Front Cell Neurosci. 2018;12:69. doi:10.3389/fncel.2018.00069.

18. Alberts B, Johnson A, Lewis J, Raff M, Roberts K, Walter P. Molecular Biology of the Cell. 5th ed. New York: Garland Science; 2007.

19. Vonesch C, Aguet F, Vonesch JL, Unser M. The colored revolution of bioimaging. IEEE Signal Processing Magazine. 2006;23(3):20–31.

20. Peng H. Bioimage informatics: A new area of engineering biology. Bioinformatics. 2008;24(17):1827–1836.

21. Swedlow JR, Eliceiri KW. Open source bioimage informatics for cell biology. Trends in Cell Biology. 2009;19(11):656–660.

22. Rittscher J. Characterization of biological processes through automated image analysis. Annual Review of Biomedical Engineering. 2010;12:315–344.

23. Danuser G. Computer vision in cell biology. Cell. 2011;147(5):973–978.

24. Meijering E. Cell segmentation: 50 years down the road [life sciences]. IEEE signal processing magazine. 2012;29(5):140–145.

25. Bengtsson E, Luengo C, Parkkinen JS, Lindblad JKT. Robust cell image segmentation methods. Pattern Recognition and Image Analysis. 2004;14(2):157–167.

26. Vincent L, Soille P. Watersheds in digital spaces: an efficient algorithm based on immersion simulations. IEEE Transactions on Pattern Analysis & Machine Intelligence. 1991;13(06):583–598.

27. Girshovitz P, Shaked NT. Generalized cell morphological parameters based on interferometric phase microscopy and their application to cell life cycle characterization. Biomedical optics express. 2012;3(8):1757–1773.

28. Burkey J. Mann-Kendall Tau-b with Sen’s Method (enhanced); 2023. https://www.mathworks.com/matlabcentral/fileexchange/11190-mann-kendall-tau-b-with-sen-s-method-enhanced.

29. Masters BR. Quantitative Phase Imaging of Cells and Tissues. Journal of Biomedical Optics. 2012;doi:10.1117/1.jbo.17.2.029901.

30. Khakh BS, Sofroniew MV. Diversity of Astrocyte Functions and Phenotypes in Neural Circuits. Nature Neuroscience. 2015;doi:10.1038/nn.4043.

31. Angermueller C, Pärnamaa T, Parts L. Deep learning for computational biology. Molecular Systems Biology. 2016;12(7). doi:10.15252/msb.20156651.

32. Lundervold A, Lundervold A. An overview of deep learning in medical imaging focusing on MRI. Zeitschrift Für Medizinische Physik. 2019;29(2). doi:10.1016/j.zemedi.2018.11.002.

